# Chimeric 3D-gastruloids – a versatile tool for studies of mammalian peri-gastrulation development

**DOI:** 10.1101/2022.05.25.493377

**Authors:** Alexandra E. Wehmeyer, Katrin M. Schüle, Alexandra Conrad, Chiara M. Schröder, Simone Probst, Sebastian J. Arnold

## Abstract

Stem cell-derived 3D-gastruloids show a remarkable capacity of self-organisation and recapitulate many aspects of gastrulation stage mammalian development. Gastruloids can be rapidly generated and offer several experimental advantages, such as scalability, observability, and accessibility for manipulation. Here, we present approaches to further expand the experimental potency of murine 3D-gastruloids by utilizing functional genetics in mouse embryonic stem cells (mESCs) to generate chimeric gastruloids. In chimeric gastruloids fluorescently labelled cells of different genotypes harbouring inducible gene-expression, or loss-of-function alleles, are combined with wildtype cells. We showcase this experimental approach in chimeric gastruloids of mESCs carrying homozygous deletions of the Tbx transcription factors *Brachyury*, or inducible expression of *Eomes*. Resulting chimeric gastruloids recapitulate reported *Eomes* and *Brachyury* functions, such as instructing cardiac fate and promoting posterior axial extension, respectively. Additionally, chimeric gastruloids revealed previously unrecognized phenotypes such as tissue sorting preference of *Brachyury*-deficient cells to endoderm, and cell non-autonomous effects of *Brachyury*-deficiency on *Wnt3a*-patterning along the embryonic axis, demonstrating some of the advantages of chimeric gastruloids as efficient tool for studies of mammalian gastrulation.

## Introduction

The versatile experimental opportunities offered by functional genetics available in the mouse as main mammalian model system have greatly enhanced our understanding of embryonic development. However, studies of mammalian embryogenesis are hampered by the relative inaccessibility of the embryo due to intrauterine growth. *Ex vivo* embryo cultures partially overcome some of the experimental restrictions but rely on the isolation of often limiting numbers of embryos. More recently, novel approaches were developed to generate stem cell derived, 3D-embryoids reflecting different stages of mammalian development from the formation of the blastocyst (Kagawa et al., 2022; Li et al., 2019; Yu et al., 2021), to periimplantation embryos (Amadei et al., 2021; Harrison et al., 2017; Harrison et al., 2018), and postimplantation stages recapitulating early organogenesis (Beccari et al., 2018; Moris et al., 2020; Turner et al., 2017; van den Brink et al., 2020; van den Brink et al., 2014; Veenvliet et al., 2020). These embryoid models demonstrate the remarkable self-organisation and robustness of embryonic programs that guide the processes of morphogenesis, growth, cell lineage specification and differentiation in the embryo as well as by *in vitro* models.

Among widely used embryoid model system are 3D-gastruloids, offering a simple experimental procedure to reproducibly generate embryoids that recapitulate development stages from early gastrulation to somitogenesis and onset of organogenesis (comparable to Embryonic days 6.5 (E6.5) – E9.0) reviewed in (van den Brink and van Oudenaarden, 2021; Veenvliet et al., 2021). 3D-gastruloids are formed by the aggregation of 100-300 mouse or human pluripotent embryonic stem cells (m or hESCs) that are treated by a 24 h pulse of the GSK3b-inhibitor CHIR to activate the canonical Wnt-cascade. This signalling stimulus induces an anterior-posterior asymmetry in the aggregate (van den Brink et al., 2014), indicated by the one-sided expression of the Tbx factor *Brachyury* that in the embryo marks the site of primitive streak formation at gastrulation onset (Rivera-Perez and Magnuson, 2005; reviewed in Arnold and Robertson, 2009). The *Brachyury*-expressing posterior pole of 3D-gastruloids elongates over several days thereby forming different tissues resembling paraxial mesoderm, neural tube, and the primitive gut tube. 3D-gastruloids show a remarkable self-organizing capacity as the different tissues are generated in proper spatial organization and arranged according to the embryonic axes (Beccari et al., 2018). 3D-gastruloids thus can be used as model systems for studies of various gastrulation-associated morphogenetic processes, as exemplified for somitogenesis (van den Brink et al., 2020; Veenvliet et al., 2020). However, some tissues of gastrulation stage embryos are less represented in 3D-organoids, including anterior mesoderm derivatives, such as cardiogenic mesoderm, and cranial structures such as cranial neural tissues (van den Brink et al., 2020; Veenvliet et al., 2020). This underrepresentation of anterior/cranial tissues most likely results from the initial induction of ESC aggregates with CHIR which imposes a strong signal for tissue identities of the posterior/caudal primitive streak (Dunty et al., 2008). However, this and other aberrations of 3D-gastruloids compared to embryos can also be used to investigate which additional regulatory requirements need to be met for proper morphogenesis of tissues and organ *anlagen* to more closely recapitulate the embryo (Veenvliet et al., 2021). Due to the scalability of this ESC-derived embryoid culture system, multiple different environmental cues can be readily tested. It is thus expected that further refinements of current protocols will lead to the generation of gastruloids that more closely resemble the full spectrum of gastrulation and following stages of embryogenesis.

Cell specification to mesoderm and definitive endoderm (DE) cell lineages is regulated by two Tbx transcription factors *Eomesodermin* (*Eomes*) and *Brachyury (T)*. Functions of both factors were previously extensively studied in mouse establishing crucial roles of *Eomes* for cell lineage specification of definitive endoderm (DE) (Arnold et al., 2008; Teo et al., 2011) and anterior mesoderm (Costello et al., 2011), and functions of *Brachyury* for the generation of posterior mesoderm derivatives, and notochord (Wymeersch et al., 2021). The compound genetic deletion of *Eomes* and *Brachyury* completely abrogates specification of any ME during differentiation of pluripotent cells (Tosic et al., 2019). While functions of these Tbx factors for specification of cell fate are well described, their roles in tissue-wide morphogenetic processes are less defined. For example, the genetic deletion of *Eomes* in the epiblast abrogates formation of the mesoderm (and DE) cell layer, hindering studies of morphogenetic functions of *Eomes*-regulated programs (Arnold et al., 2008). Similarly, studies of the cell-intrinsic roles of *Brachyury* in posterior body axis extension are compromised by cell non-autonomous functions of *Brachyury* in feed-forward regulatory loops to maintain caudal Wnt-signals, that could independently act on cell specification and morphogenetic programs (Arnold et al., 2000; Dunty et al., 2008; Martin and Kimelman, 2008; Turner et al., 2014; Yamaguchi et al., 1999). These constraints to distinguish cell-autonomous effects of loss-of-gene-function from secondary effects, such as by general disturbance of morphogenesis, can be experimentally addressed by the generation of embryonic chimeras. Here, cell-autonomous gene functions can be studied when limited numbers of genetically altered cells are placed in an otherwise wildtype cellular environment. Chimeric approaches were previously reported for studies of *Brachyury* functions, such as in zebrafish (Halpern et al., 1993) and mouse (Wilson et al., 1993; Wilson et al., 1995; Wilson and Beddington, 1997), as well as for *Eomes* functions (Arnold et al., 2008), indicating regulatory roles of both Tbx factors in cell migration and morphogenesis. However, embryonic chimera analyses are highly labour and resource intensive, especially in mouse, and thus are only infrequently employed.

In this report, we demonstrate an approach to further increase the experimental potency of murine 3D-gastruloids by expanding it to chimeric analyses. We use genetically engineered, traceable mESCs to generate chimeric 3D-gastruloids that are composed of cells with different genotypes. To showcase the advantages of this approach and to test experimental feasibility we generated fluorescently labelled mESC lines with homozygous loss-of-function, or inducible expression cassettes for the two Tbx transcription factors *Brachyury* and *Eomes*. These are used to generate chimeric gastruloids, e.g. by mixing defined ratios of inducible Tbx-expressing, or Tbx-deficient and WT cells to generate chimeric situations with defined cell contribution that are normally only difficult to achieve by conventional genetic tools or by cell injections into embryos. In addition to demonstrating the efficiency and feasibility of this experimental approach of chimeric gastruloids, this study also provides some previously unrecognized insights into the morphogenetic functions of *Eomes* and *Brachyury*.

## Results and Discussion

To expand the experimental versatility of the 3D-gastruloid model system we combined it with the use of genetically modified mESCs and fluorescent imaging. We generated chimeric gastruloids composed of cells with different genetic backgrounds that can be readily traced by different fluorescent membrane labels (Fig. 1A, B). WT mESCs are permanently labelled by monoallelic knock-in of membrane GFP (mG) into the Rosa26 genomic locus (Fig. 1C), while membrane Tomato (mT) labelling was used for genetically manipulated mESCs lines (Fig. 1C). Genetic alterations used in presented experiments were the homozygous deletion of the Tbx transcription factor *Brachyury* (*Bra*^*-/-*^), and the ICE-mediated (induced cassette exchange) targeted integration of *Eomes* 3’ to a monoallelic Tet-responsive element (TRE) in A2lox (E14-based) mESCs (Iacovino et al., 2014), resulting in TRE.*Eomes*GFP (short TRE.*Eo*). These cell lines were first tested for their abilities to form regular gastruloids. This demonstrated that fluorescent membrane labels don’t have an impact on gastruloid formation, while *Bra*^*-/-*^ and dox-induced TRE.*Eo* cells show a failure to form regularly extended gastruloids at 120 h (Supplementary Fig. 1). To generate chimeric gastruloids we followed two approaches (Fig. 1A, D, E), either by mixing of different cells during the aggregation of mESCs at the beginning of gastruloid culture (Fig. 1E), or by merging of two preformed mESCs aggregates composed of different cells 24 h after the initial aggregation (Fig. 1 B, D). Such merged ESC aggregates rapidly fuse and stably adhere after placing them together in 96-wells (Fig. 1D, Supplementary Movie 1) or in hanging drops (not shown).

**Fig. 1.**
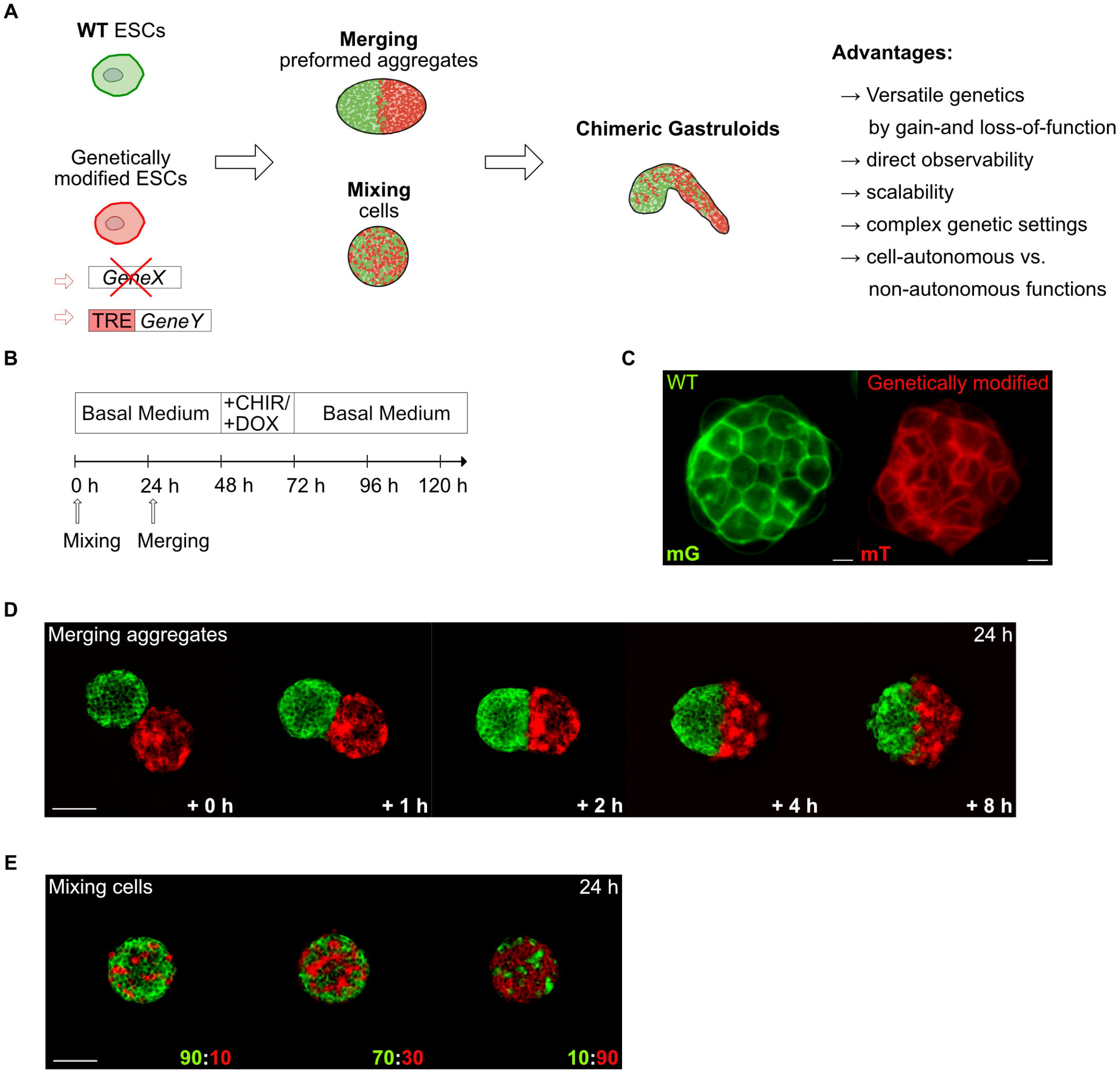
Experimental approaches for the generation of chimeric gastruloids. **(A)** Schematic of two alternative experimental approaches to generate chimeric gastruloids from fluorescently labelled embryonic stem cells (mESCs). Wildtype (WT) mESCs are marked by membrane-GFP (mG) and combined with membrane-Tomato (mT) labelled, genetically modified mESCs. The genetic modifications of mESCs comprise homozygous gene deletions (*GeneX*) or inducible gene-expression by the targeting of cDNAs into a fully controllable pre-engineered locus containing a Tetracycline responsive element (TRE) for dox-dependent induction of gene expression (*GeneY*). Chimeric gastruloids are generated by either mixing of mESCs at the beginning of gastruloid culture, or by merging preformed mESCs aggregates preceeding the induction of gastruloids by CHIR. Chimeric gastruloids can be used as versatile novel model system for various types of studies and embryonic research questions as indicated exemplary. **(B)** Schematic of the protocol to form chimeric gastruloids indicating timepoints for either mixing of cells, or merging of aggregates to create chimeric gastruloids. **(C)** Membrane-GFP (mG) labelling of WT mESCs and membrane-Tomato (mT) labelling for genetically modified mESCs allows for distinguishing different cell types in chimeric gastruloids. Scale bars 10 μm. **(D)** Timelapse-imaging of the merging process of pre-formed ESC aggregates at indicated time points. 150 mG and mT mESCs were aggregated for 24 h before merging them by placing them together in 96-well plates. After 1 hr two aggregates spontaneously aggregated and formed stable contacts. Scale bars 100 μm. Also see **Supplementary Movie 1** for a 12 h time-lapse movie. **(E)** Examples of mixed ESC aggregates at 24 h after aggregation of 300 mESCs by mixing mG WT mESCs and mT *Bra*^*-/-*^ mESCs at indicated ratios. Scale bars 100 μm.

The mixing of cells at different ratios can be used to evaluate cell-autonomous vs. cell non-autonomous gene functions, by testing how different levels of cell contribution affects tissue behaviour, e.g. when using cells with loss-of-gene-function. Merging of cell aggregates can be applied when the behaviour of coherent groups of cells is analysed, such as in studies of inductive tissue interactions. A similar approach was recently reported where a small aggregate of 50 cells was treated with BMP4 to induce organizer-activities which was used instead of CHIR-treatment for the induction of gastruloid-formation (Xu et al., 2021). Thus, chimeric gastruloids offer multiple experimental opportunities to study gastrulation development (Fig. 1A) as exemplarily demonstrated in following experiments.

First, we tested if chimeric gastruloids are a suitable model to study instructive gene functions during lineage specification. Hence, we generated chimeric gastruloids by merging preformed aggregates of WT mESCs (mG-label) and mESCs containing the doxycycline (Dox) inducible *Eomes*.GFP expression cassette (TRE.*Eo*; mT-label) (Fig. 2A, B). *Eome*s is crucially required for lineage specification towards definitive endoderm (Arnold et al., 2008; Teo et al., 2011) and anterior mesoderm, including the cardiac lineage (Costello et al., 2011; Probst et al., 2021). We tested if forced *Eomes* expression in a group of cells (TRE.*Eo*) within a gastruloid would be sufficient to induce a coherent heart-like domain, which is only inconsistently forming in gastruloids composed of WT mESCs (Rossi et al., 2021; van den Brink and van Oudenaarden, 2021). Indeed, Dox-induced *EomesGFP*-expressing cells form a domain of beating cardiomyocytes (Fig. 2C, Supplementary movie 2) in chimeric gastruloids in 33.7% (± 3.0 SD) of cases, whereas similar beating areas are only observed in 1.5% (± 2.5 SD) in uninduced chimeric gastruloids (Fig. 2D) or using only A2lox WT mESCs (not shown). To correlate induced *EomesGFP* expression with the cardiogenic domain we performed *in situ* hybridization analyses for early cardiac markers *Mlc2a* and *Nkx2*.*5*, which are expressed in mT-marked TRE.*Eo* cells that are forming a coherent domain on one side of gastruloids (Fig. 2E, F). To analyse the position of TRE.*Eo* cell-derived domains along the AP axis of merged chimeric gastruloids, we performed immunostainings against BRACHYURY and CDX2, indicating the posterior pole of extending gastruloids (Fig. 2G, H). The domains of TRE.*Eo* cells are predominantly found on one side along the AP-axis of the gastruloids and oriented towards the anterior pole (Fig. 2G, H). Of note, these coherent domains of cells are not found in gastruloids when two preformed aggregates of WT cells are merged where cells undergo increased mixing (Supplementary Fig. 2A). Interestingly, TRE.*Eo* cell containing gastruloids consistently showed a reduction in axis extension, so that the anterior-posterior (AP) axis of resulting gastruloids is less obvious in comparison to only WT-derived gastruloids (Supplementary Fig. 2B, C). In short, these experiments show that induced expression of *Eomes* suffices to cell-autonomously generate coherent cardiogenic domains oriented to the anterior-lateral side of developing gastruloids in a highly increased proportion of gastruloids compared to uninduced gastruloids.

**Fig. 2.**
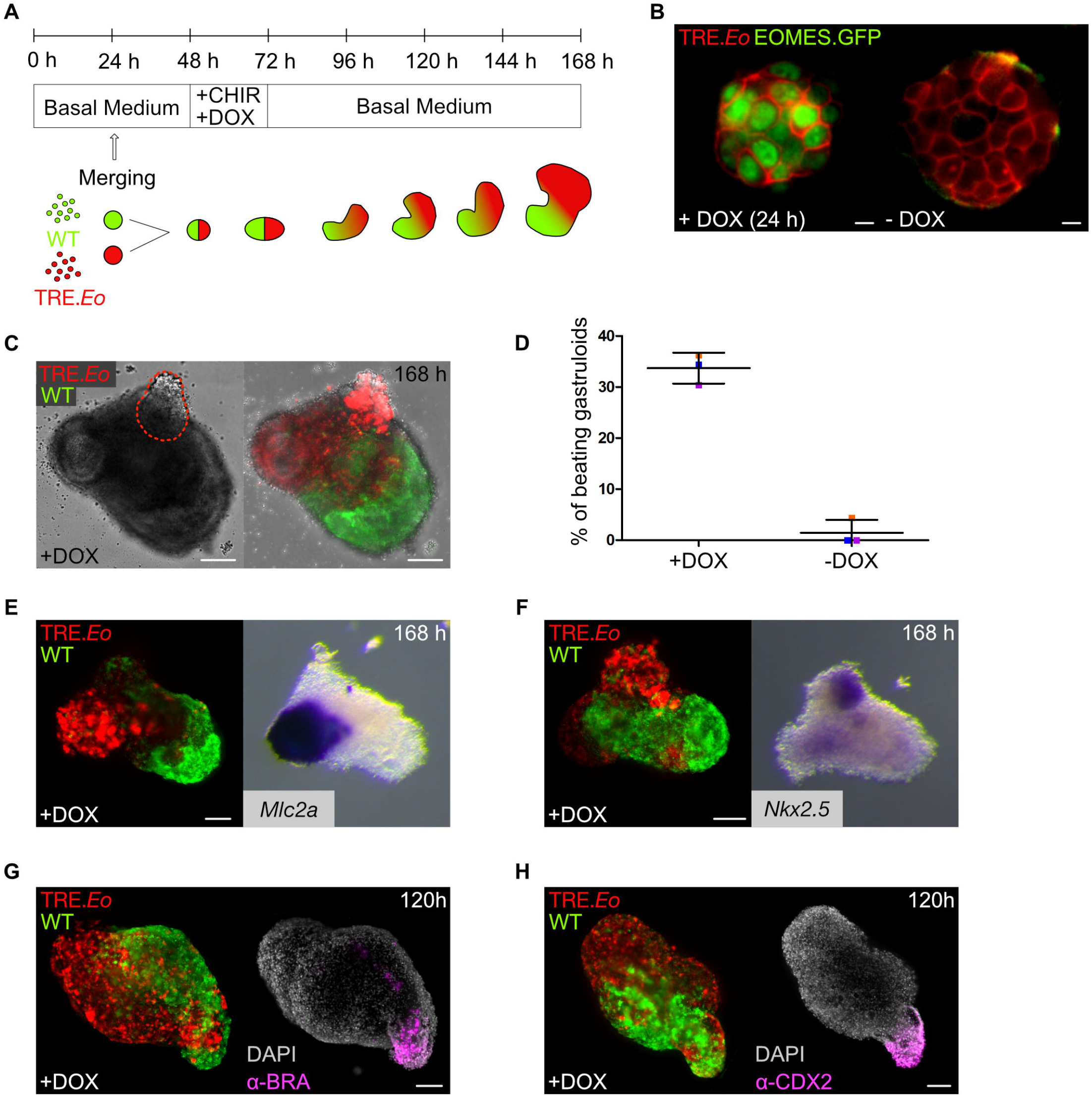
Instructive functions of *Eomes* for cardiac lineage specification in merged chimeric gastruloids. **(A)** Schematic illustrating the generation and culture of chimeric gastruloids by merging preformed aggregates of mG-labelled WT mESCs and mT-labelled mESCs harbouring *EomesGFP* in the Doxycycline (Dox)-inducible gene locus (TRE.*EomesGFP, short* TRE.*Eo*). **(B)** Fluorescent microscopy of TRE.*Eo* mESCs showing nuclear expression of doxycycline-induced (+DOX) EOMES.GFP after 24 h of administration that is absent in -DOX conditions. Scale bars 10 μm. **(C)** Brightfield (left) and fluorescent (right) images of a chimeric gastruloid at 168 h following induced *Eomes* expression showing the region of beating cardiomyocytes within the gastruloid that is mostly derived of mT-labelled TRE.*Eo* cells. The dashed line indicates the domain of beating cells. See also **Supplementary Movie 2** for the corresponding time-lapse movie. **(D)** Statistics comparing the percentage of beating gastruloids at 168 h in gastruloids with forced *Eomes* expression +DOX (33.7% ± 3.0) and uninduced controls -DOX (1.5% ± 2.5) in n=3 independent experiments that are colour-encoded in orange, blue, and purple. Error bars indicate SD. **(E, F)** Fluorescent microscopy (left) and whole-mount *in situ* hybridization (right) of the same chimeric gastruloids following induced *Eomes*-expression (+DOX) showing instructive functions of forced *Eomes*-expression in TRE.*Eo* cells for the induction of cardiac progenitors, marked by expression of (**E**) *Mlc2a* (n=12), and (**F)** *Nkx2*.*5* (n=10). **(G, H)** Fluorescent microscopy (left) and immunofluorescence staining (right) showing positioning of TRE.*Eo* cells at the opposite pole of posterior **(G)** BRACHYURY (a-BRA) (n=24) and **(H)** CDX2 (n=10) expression. Scale bars 100 μm in C, E-H.

Next, we aimed for testing the feasibility of chimeric gastruloids for the assessment of cell-autonomous vs. cell non-autonomous functions of *Brachyury* during posterior elongation, as the shortening of the posterior body axis is the most prominent phenotype of *Brachyury*-mutant embryos (Fig. 3A)(Wymeersch et al., 2021). Gastruloids entirely generated from mT-labelled *Bra*^-/-^ mESCs present with a phenotype of impaired elongation, reminiscent of the failure of tailbud elongation observed in *Brachyury*-mutant embryos (Fig. 3B, E; Supplementary Figs. 1, 3B) (Inman and Downs, 2006). Increasing the proportional contribution of WT cells (mG) to the gastruloid by intermixing WT with *Bra*^*-/-*^ mESCs during the initial aggregation of 300 mESCs leads to the gradual extension and increase in overall tissue mass of the posterior portion of the mixed gastruloids (Fig. 3B). We analysed BRACHYURY presence in the WT cells in the posterior regions of chimeric gastruloids with high (80%) and equal (50%) contribution of *Bra*^*-/-*^ cells at 24 h time intervals. The posterior region of chimeric gastruloids with a contribution of *Bra*^*-/-*^ cells of 80% show less confined and reduced a-BRACHYURY staining (Fig. 3C) compared to chimeric gastruloids where *Bra*^*-/-*^ cell contribution is 50%. Here, *Brachyury* expression is found robustly but with decreased intensity in the posterior pole when compared to WT gastruloids (Fig. 3D, and Supplementary Fig. 3A).

**Fig. 3.**
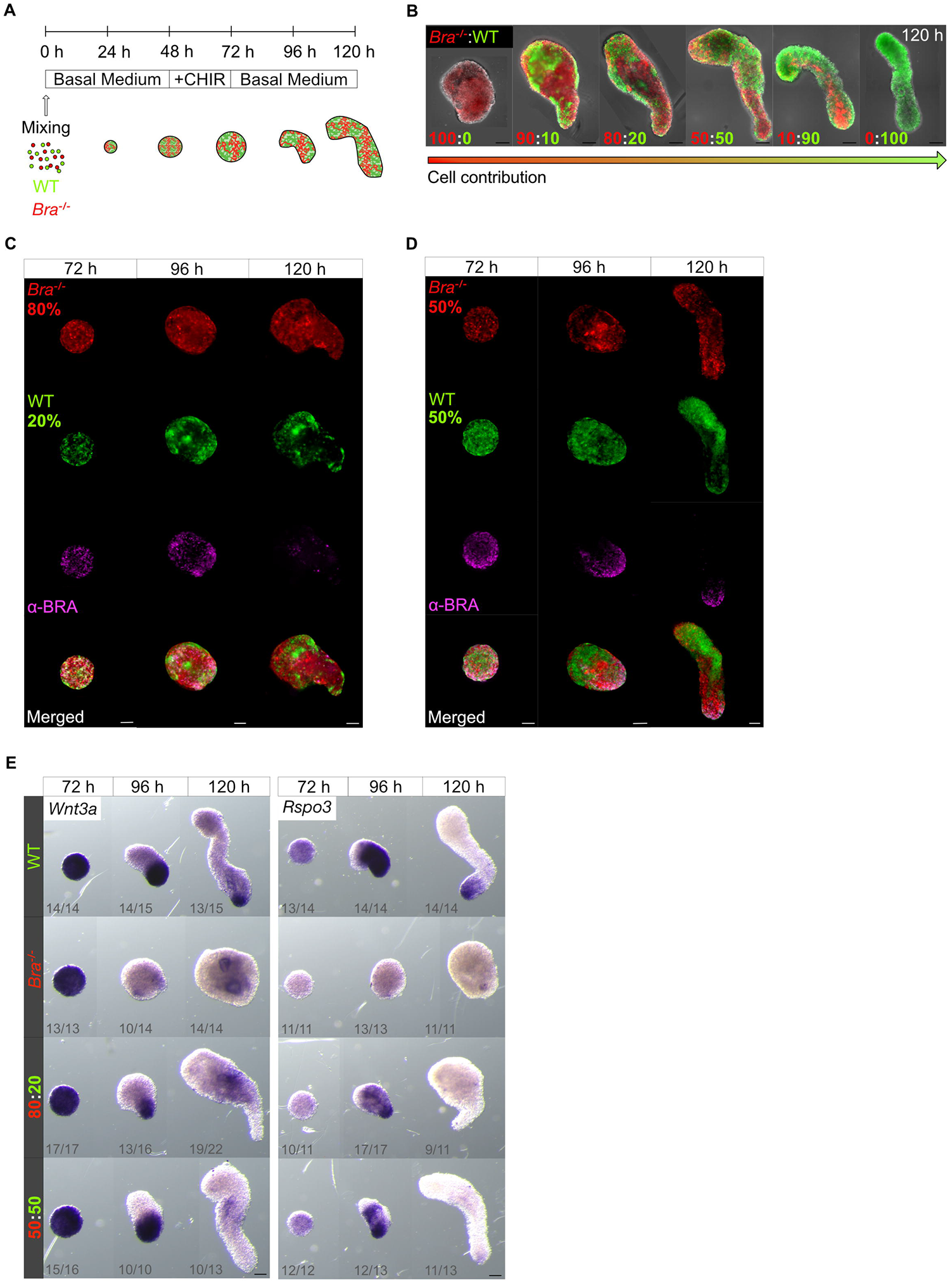
*Brachyury* functions during axis-elongation in chimeric gastruloids generated by cell mixing with high *Bra*^*-/-*^ cell contribution. **(A)** Schematic illustrating the generation of chimeric gastruloids by mixing of mG-labelled WT mESCs and mT-labelled Brachyury-deficient (*Bra*^*-/-*^) mESCs. **(B)** Chimeric gastruloids at 120 h generated by mixing of *Bra*^*-/-*^ (mT) and WT (mG) mESCs at indicated ratios of cell numbers. Gastruloids with a contribution of *Bra*^*-/-*^ cells above 80 % show reduced axial elongation. Gastruloids entirely generated from *Bra*^*-/-*^ cells fail to axially extend beyond an oval shape. At WT (mG) cell contribution > 50 % axial elongation is similar to WT gastruloids but occasionally shows a thinning of the posterior portion. Across all experiments *Bra*^*-/-*^ (mT) cells preferentially contribute to the posterior portion of mixed gastruloids. Anterior is oriented to the top and posterior to the bottom in all images. **(C, D)** Immunofluorescence staining against BRACHYURY (a-BRA) in gastruloids generated by cell mixing of *Bra*^*-/-*^ and WT cells in ratios of 80:20 (**C**) and 50:50 (**D**) at indicated timepoints, showing less confined and reduced BRACHYURY expression in 80:20 chimeras at 96 and 120 h in comparison to posteriorly localized, robust BRACHYURY expression in 50:50 chimeras. Images are representatives of n=14 (72 h); n=9 (96 h); n=14 (120 h) in (**C**), and n=16 (72 h); n=25 (96 h), n=18 (120 h) gastruloids in (**D**). **(E)** Whole-mount *in situ* hybridization for *Wnt3a* and *Rspo3* of WT, Bra^-/-^-null and chimeric gastruloids of indicated cell ratios at 72 h, 96 h, and 120 h showing mislocalised and reduced expression of *Wnt3a* and *Rspo3* in Bra-deficient and chimeric WT: Bra^-/-^ mixed gastruloids. The frequencies of representative staining pattern per numbers of examined gastruloid are indicated below each image. Scale bars 100 μm in B-D.

The absence of *Brachyury* expression also from WT cells in gastruloids with high contribution of *Bra*^*-/-*^ cells suggests cell non-autonomous effects of *Bra*^*-/-*^ cells. These are most likely explained by disturbances of the previously described feed-forward regulation of *Wnt3a* and *Brachyury* in the tail-bud region (Arnold et al., 2000; Dunty et al., 2008; Martin and Kimelman, 2008; Turner et al., 2014; Yamaguchi et al., 1999). To test this, we performed *in situ* hybridization to analyse expression of *Wnt3a* and the putative *Brachyury* target gene and Wnt co-ligand *Rspo3 (*Koch *et al*., 2017) in WT gastruloids, in gastruloids entirely composed of *Bra*^*-/-*^ cells, and in mixed gastruloids with 80% and 50% *Bra*^*-/-*^ cell contribution at 72, 96 and 120 h (Fig. 3E). This analysis demonstrates *Brachyury-*requirements for the sustained expression of *Wnt3a*, which is absent from *Bra*^*-/-*^ gastruloids at 96 h and greatly reduced and mislocalised in mixed gastruloids at 120 h. Interestingly *Wnt3a* expression, which normally is confined to the posterior pole, is found at intermediate levels of mixed gastruloids, most likely resulting from unequal cell distribution of WT and *Bra*^*-/-*^ cells, which accumulate at the posterior of mixed *Bra*^*-/-*^: WT gastruloids (see also Fig. 4 and discussion below). Analysis of *Rspo3* expression recapitulates the lack of *Brachyury* expression in *Bra*^*-/-*^ gastruloids and the prematurely exhausted *Brachyury* expression in *Bra*^*-/-*^: WT mixed gastruloids (Fig. 3C-E).

**Fig. 4.**
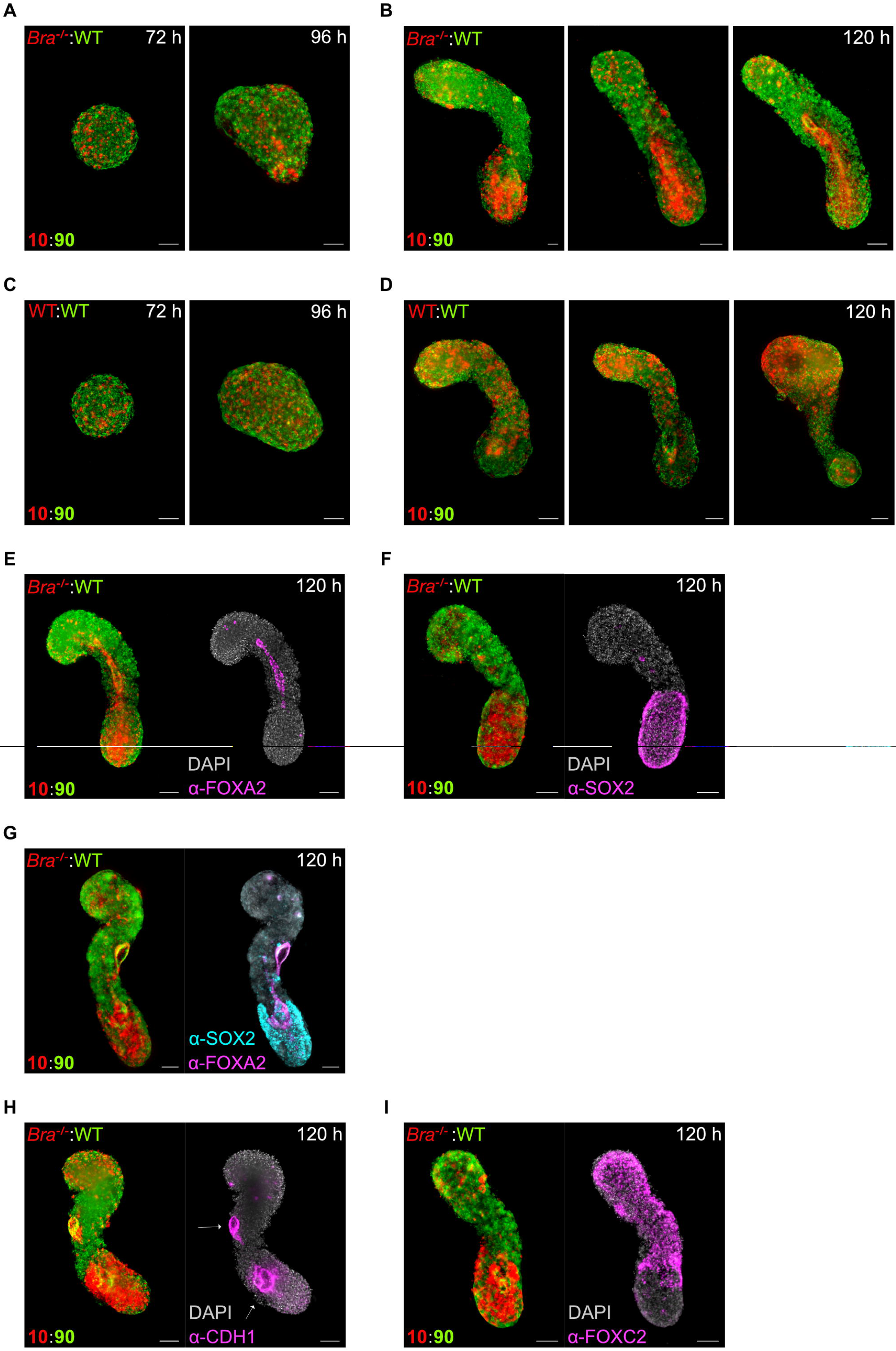
Brachyury functions for cell lineage specification and tissue sorting in chimeric gastruloids with low *Bra*^*-/-*^ cell contribution. **(A-D)** Chimeric gastruloids generated by mixing of 10% *Bra*^*-/-*^ (mT) and 90 % WT (mG) ESCs at **(A)** 72 h and 96 h, and **(B)** three representative gastruloids at 120 h showing the spectrum of distribution of *Bra*^*-/-*^ cells between the posterior pole and the midline of chimeric gastruloids. (**C, D**) Control chimeric gastruloids generated by mixing of 10% WT (mT) and 90% WT (mG) ESCs at **(C)** 72 h, 96 h, and **(D)** three replicates at 120 h don’t show distinct patterns of cell distribution. **(E-G)** Immunofluorescence staining against (**E**) FOXA2 (n=6) and (**F**) SOX2 (n=8) individually, and **(G)** double IF staining showing the tissue contribution of *Bra*^*-/-*^ cells to FOXA2+ endoderm and SOX2+ posterior cells (n=14). **(H)** *Bra*^*-/-*^ cells contribute to the forming primitive gut tubes (arrows) and are positive for the epithelial marker CDH1 (E-Cadherin) (n=3). **(I)** *Bra*^*-/-*^ cells are mostly excluded from a mesodermal domain that is positive for FOXC2 (n=7). Scale bars 100 μm.

Next, in addition to the tissue-wide, partially cell non-autonomous phenotype of *Brachyury*-deficiency in gastruloids with a high contribution of *Bra*^-/-^ cells, we analysed the cell-autonomous effects of *Brachyury*-deficiency in conditions with low contribution of mutant cells (Fig. 4). We generated chimeric gastruloids by mixing cells in a 10:90 ratio of *Bra*^*-/-*^: WT mESCs (Fig. 4A, B) and compared resulting gastruloids with controls where 10% mT-labelled WT cells were used (Fig. 4C, D). Further controls were included where cells were mixed at 50:50 ratios (Supplementary Fig. 4A, B). WT cells randomly disperse, while *Bra*^*-/-*^ exhibit a tissue sorting behaviour (Fig. 4A-D, Supplementary Fig. 4A, B). At 120 h of culture *Bra*^*-/-*^ cells are predominantly found along the midline in the interior of the gastruloids, and in the posterior pole (Fig. 4B). To determine tissue identity of these regions we used immunofluorescence (IF) staining against FOXA2 and CDH1 (E-Cadherin), labelling DE (Viotti et al., 2014), SOX2 labelling neural tube progenitors (Wood and Episkopou, 1999), and FOXC2 as mesoderm marker, and combined IF-staining with the fluorescent labels of WT (mG) and *Bra*^*-/-*^ cells (mT) (Fig. 4E-I). FOXA2 and CDH1 stainings indicate that *Bra*^*-/-*^ cells are biased towards DE and favored to form the gut tube-like structure in the midline of gastruloids (Fig. 4E, G, H). This finding not only demonstrates that *Brachyury* is dispensable for DE lineage specification from pluripotent cells, but suggests that *Brachyury* actually counteracts DE specification programs and thus may not represent a suitable marker for early DE-forming cells as previously suggested (Kubo et al., 2004). In addition to their contribution to endoderm structures of chimeric gastruloids, *Bra*^*-/-*^ cells are predominantly found in the posterior region of chimeric gastruloids. Here, *Bra*^*-/-*^ cells show SOX2 staining marking them as neural tube progenitors (Fig. 4F, G). Furthermore, *Bra*^*-/-*^ cells are rather excluded from the FOXC2-positive mesoderm forming domain (Fig. 4I), in accordance with previous findings about NE-repressive and mesoderm promoting functions of *Brachyury* (Koch et al., 2017; Tosic et al., 2019).

In conclusion, this study illustrates the feasibility and experimental potency to use functional genetics in mESCs to generate chimeric 3D-gastruloids. These allow for the rapid and reproducible analysis of gain- and loss-of-gene functions, which may reveal different phenotypic outcomes according to the degree of cellular contribution in chimeric gastruloids. Close attention should be paid to control for the general variability in mESC morphogenetic and differentiation behaviour, which might not strictly depend on the genetic manipulation of each cell line. We did not experience these problems in presented experiments, as they were all based on usage of mESCs lines originating from the same parental clone (A2lox, Iacovino et al., 2014).

Here, we used the inducible expression of the Tbx transcription factor *Eomes* to demonstrate how limitations of 3D gastruloids can be overcome by genetically providing regulatory cues that are missing or underrepresented in CHIR-only treated gastruloids, namely the induced formation of anterior mesoderm derivatives including heart tissue. The analysis of *Brachyury*-deficient cells in chimeric gastruloids remarkably reflects aspects previous studies of embryonic chimeras (Wilson et al., 1993; Wilson et al., 1995; Wilson and Beddington, 1997). Here, *Bra*^*-/-*^ cells progressively accumulate in the tailbud region, most likely as result of *Brachyury* functions in regulation of mesoderm cell migration leading to passive replacement of mutant cells into the most posterior embryonic regions (Wilson et al., 1995; Wilson and Beddington, 1997). It thus will be interesting to follow morphogenetic cell movements of *Bra*^*-/-*^ cells in chimeric gastruloids. In addition, the analysis of *Brachyury* deficient cells in chimeras with low cell contribution showed a cell-autonomous bias towards the DE, a cell lineage that strictly depends on *Eomes* functions (Arnold et al., 2008; Teo et al., 2011). Since *Brachyury* and *Eomes* are co-expressed in cells of the early primitive streak in early gastrulating embryos (Tosic et al., 2019), this poses the interesting question about reciprocal regulatory interactions between these two related Tbx factors.

In summary, this study demonstrates that chimeric gastruloids represent a powerful experimental tool for the analysis of gastrulation stage embryogenesis. Gastruloids already exhibit a high degree of experimental accessibility, observability and scalability. The additional implementation of chimeric gastruloids allows to analyse gene-functions at cellular level without affecting the general gastruloid architecture and might thus enhance our understanding of cell-signal interactions during gastrulation morphogenesis.

## Materials and Methods

### Cell lines

A2lox mESCs (Iacovino et al., 2014) were cultured in Dulbecco’s modified Eagle’s medium (DMEM) containing 15 % fetal bovine serum (FBS, Gibco), 2 mM L-glutamine, 1X non-essential amino acids (NEAA), 1 mM sodium-pyruvate, 1X penicillin/streptomycin (all from Gibco), 100 μM β-mercaptoethanol (Sigma), Leukemia inhibitory factor (ESGRO LIF, Merck Millipore, 1000 U/ml), and 2i: CHIR99021 (Axon Medchem, 3 μM) and PD0325901 (Axon Medchem, 1 μM) on 0.1% gelatine-coated dishes. The medium was changed daily and mESCs were passaged every other day. The generation of homozygous *Bra*^*-/-*^ mESCs and of A2lox mESCs harbouring monoallelic dox-inducible expression cassette for *Eomes*.GFP were described previously (Tosic et al., 2019). A2lox cells with membrane-tagged fluorescent labels (membrane-Tomato, mT, and membrane-GFP, mG) were generated by targeted integration of a mT/mG targeting vector (Muzumdar et al., 2007), Addgene plasmid #17787) into the Rosa26 locus. 1x10^6^ A2lox mESCs (WT and *Bra*^*-/-*^) were transfected with 2.5 μg of linearized vector using the Nucleofector ESC kit (Lonza) and G418 selected (350 μg/ml) on a monolayer of MitoC (Sigma)-mitotically inactivated STO feeder cells. mT-expressing ESC clones were picked on day 9 of selection. To convert the expression of the membrane-Tomato (mT) to membrane-GFP (mG) in WT A2lox mESCs, cells were treated for 24 h with 5 μg/ml Doxycyclin (Sigma, D9891) for induced expression of the Cre-recombinase from the Dox-inducible locus of A2lox WT cells to excise the loxP-flanked mT expression cassette and bring the mG expression cassette under the transcriptional control of the Rosa26 gene locus. After Cre-excision mG-expressing WT A2lox mESCs underwent one round of clonal selection by minimal dilution of 500 cells onto a 10 cm cell culture dish.

### Generation of chimeric gastruloids

Gastruloids were generated using published protocols (van den Brink et al., 2020) with some modifications as outlined below. Gastruloid formation was performed in ESGRO Complete Basal Medium (Merck Millipore) in the absence of Matrigel. To generate chimeric gastruloids using different mESCs by merging of preformed aggregates 150 cells of each ESC line were aggregated in 40 μl of ESGRO basal medium in 96-well format (Greiner ultra-low attachment plates, No. 650970) for 24 h before merging by combining two aggregates into the same well. At 48 h following first aggregate formation, fused gastruloids were induced by administration of 3 μM CHIR and if indicated with doxycycline (1 μg/μl, Sigma) for 24 h. In the course of gastruloid culture, the medium was changed daily at 72, 96, 120 and 144 h. For the generation of gastruloids by mixing of different ESC lines, aggregates were formed from a total of 300 mESCs at various ratios between the two ESC lines, and further gastruloid formation followed by previously used protocols (van den Brink et al., 2020).

### Whole mount *in situ* hybridization

Whole-mount *in situ* hybridization was performed according to standard protocols using standard probes for *Mlc2a, Nkx2*.*5, Wnt3a*, and *Rspo3* (Details can be requested from the authors). In brief, gastruloids were fixed in 4% PFA/PBS o/n at 4°C, dehydrated via a methanol series and stored in methanol at -20°C. After rehydration gastruloids were bleached in 6 % H_2_O_2_ for 5 min, digested by 1.6 μg/ml ProteinaseK in PBT for 2 min, and postfixed in 4% PFA/0,2% glutaraldehyde for 20 min before prehybridization for 2 h and hybridization o/n according to standard protocols. DIG-labelled RNA probe was detected using anti-Digoxigenin-AP Fab fragments (Roche) in 1% sheep serum, 2% BBR in MAB (0.1 M Maleic acid, 0.3 M NaCl, NaOH, 1% Tween-20 in H_2_O, pH 7.5) and incubation at 4°C o/n. Antibody was washed out by extensive washes in MAB (>24 h, RT), and color reaction performed in BM purple staining solution (Roche) for 2-6 h at RT.

### Whole mount immunofluorescence staining

Gastruloids were fixed in 4% PFA /PBS for 1 h at 4°C, permeabilized (0.3% Triton X-100/PBT, 30 min) and blocked in 1% BSA / PBT for 1h at RT. Primary antibody incubation was performed at 4°C o/n in 1% BSA / PBT, gastruloids washed 4x in PBT before secondary fluorescence-conjugated antibody incubation for 3 h followed by DAPI staining for 30 min at RT. Primary antibodies used were BRACHYURY (R&D Systems; AF2085), CDX2 (Biogenex; MU392A) FOXA2 (Cell Signaling; 8186S), E-Cadherin (BD Transduction Laboratories; 610182), SOX2 (R&D Systems; AF2018), FOXC2 (R&D Systems; AF6989) at suggested dilutions. Secondary anti-goat, anti-rabbit, anti-mouse and anti-sheep Alexa Fluor 647-conjugated antibodies (Thermo Fisher), and anti-goat CF405M (Biotium).

### Histological sections of gastruloids

Fixed gastruloids were processed through 15% and 30% sucrose/PBS and incubated for at least 1h in embedding medium (15% sucrose, 7,5% gelatin in PBS) before cryo-embedding. Embedded gastruloids were cut with into 8 μm sections, mounted with ProLong Diamond Antifade Mountant (Life Technologies, P36970) and imaged as described below.

### Imaging

Images were acquired on a Leica DMi8 Thunder Imager System or a Leica M165FC Stereo microscope. Images were processed in the Leica LASX software and Affinity Photo. During time lapse imaging gastruloids were maintained under constant conditions at 37°C, 5% CO_2_.

### Data acquisition and statistics

The percentage of beating gastruloids (Fig. 2D) was determined by manually counting during life imaging in three individual experiments. Following numbers of gastruloids were evaluated: 23 (+DOX) and 21 (-DOX) for n1; 61(+DOX) and 57 (-DOX) for n2, and 91(+DOX) and 90 (-DOX) for n3.

The length of the anterior-posterior axis of gastruloids (Fig. 2J) was measured using the LASX software (Leica) for n=34 (WTmG-WTmT) and n=36 (WTmG-TRE.*Eo*mT). Graphs (Scatter plot; Mean with SD) and column statistics were acquired with the GraphPad Prism software (Version 5.04).

## Acknowledgements

We thank T. Bass for excellent technical assistance and Michael Kyba for the A2lox ESC line.

## Funding

This work was supported by the German Research Foundation (DFG) through the Heisenberg Program (AR 732/3-1), project grant (AR 732/2-1,) project B07 of SFB 1140 (project ID 246781735), project A03 of SFB 850 (project ID 89986987), project P7 of SFB 1453 (project ID 431984000), and Germany’s Excellence Strategy (CIBSS – EXC-2189 – Project ID 390939984) to S.J.A., and by the MOTI-VATE program of the Freiburg Medical Faculty supported by the Else-Kröner-Fresenius-Stiftung to A.C.

## Competing interests

The authors declare no competing interests.

## Author contributions

A.E.W., K.M.S., A.C., C.M.S. generated different ESC lines and performed experiments. A.E.W, S.P. and S.J.A. planned and analysed experiments. A.E.W. and S.J.A. prepared figures, wrote and edited the manuscript with input from all authors. S.J.A. conceived the study.

## Notes

### Competing Interest Statement

The authors have declared no competing interest.

### Summary of Updates

The manuscript was comprehensively revised and now includes several new experiments and data points, so that the manuscript now encompasses four main and four supplementary figures. Mainly some basic descriptions of the experimental model is now included as well as a previously missing experiment to explain suggested cell non-autonomous effects of Brachyury-deficiency on body axis elongation via feedback loops with Wnt3a.

## References

Amadei, G., Lau, K. Y. C., De Jonghe, J., Gantner, C. W., Sozen, B., Chan, C., Zhu, M., Kyprianou, C., Hollfelder, F. and Zernicka-Goetz, M. (2021). Inducible Stem-Cell-Derived Embryos Capture Mouse Morphogenetic Events In Vitro. Dev Cell 56, 366–382 e369.

Arnold, S. J., Hofmann, U. K., Bikoff, E. K. and Robertson, E. J. (2008). Pivotal roles for eomesodermin during axis formation, epithelium-to-mesenchyme transition and endoderm specification in the mouse. Development 135, 501–511.

Arnold, S. J. and Robertson, E. J. (2009). Making a commitment: cell lineage allocation and axis patterning in the early mouse embryo. Nat Rev Mol Cell Biol 10, 91–103.

Arnold, S. J., Stappert, J., Bauer, A., Kispert, A., Herrmann, B. G. and Kemler, R. (2000). Brachyury is a target gene of the Wnt/beta-catenin signaling pathway. Mech Dev 91, 249–258.

Beccari, L., Moris, N., Girgin, M., Turner, D. A., Baillie-Johnson, P., Cossy, A. C., Lutolf, M. P., Duboule, D. and Arias, A. M. (2018). Multi-axial self-organization properties of mouse embryonic stem cells into gastruloids. Nature 562, 272–276.

Costello, I., Pimeisl, I. M., Drager, S., Bikoff, E. K., Robertson, E. J. and Arnold, S. J. (2011). The T-box transcription factor Eomesodermin acts upstream of Mesp1 to specify cardiac mesoderm during mouse gastrulation. Nat Cell Biol 13, 1084–1091.

Dunty, W. C., Jr., Biris, K. K., Chalamalasetty, R. B., Taketo, M. M., Lewandoski, M. and Yamaguchi, T. P. (2008). Wnt3a/beta-catenin signaling controls posterior body development by coordinating mesoderm formation and segmentation. Development 135, 85–94.

Halpern, M. E., Ho, R. K., Walker, C. and Kimmel, C. B. (1993). Induction of muscle pioneers and floor plate is distinguished by the zebrafish no tail mutation. Cell 75, 99–111.

Harrison, S. E., Sozen, B., Christodoulou, N., Kyprianou, C. and Zernicka-Goetz, M. (2017). Assembly of embryonic and extraembryonic stem cells to mimic embryogenesis in vitro. Science 356.

Harrison, S. E., Sozen, B. and Zernicka-Goetz, M. (2018). In vitro generation of mouse polarized embryo-like structures from embryonic and trophoblast stem cells. Nat Protoc 13, 1586–1602.

Iacovino, M., Roth, M. E. and Kyba, M. (2014). Rapid genetic modification of mouse embryonic stem cells by Inducible Cassette Exchange recombination. Methods Mol Biol 1101, 339–351.

Inman, K. E. and Downs, K. M. (2006). Brachyury is required for elongation and vasculogenesis in the murine allantois. Development 133, 2947–2959.

Kagawa, H., Javali, A., Khoei, H. H., Sommer, T. M., Sestini, G., Novatchkova, M., Scholte Op Reimer, Y., Castel, G., Bruneau, A., Maenhoudt, N., et al. (2022). Human blastoids model blastocyst development and implantation. Nature 601, 600–605.

Koch, F., Scholze, M., Wittler, L., Schifferl, D., Sudheer, S., Grote, P., Timmermann, B., Macura, K. and Herrmann, B. G. (2017). Antagonistic Activities of Sox2 and Brachyury Control the Fate Choice of Neuro-Mesodermal Progenitors. Dev Cell 42, 514–526 e517.

Kubo, A., Shinozaki, K., Shannon, J. M., Kouskoff, V., Kennedy, M., Woo, S., Fehling, H. J. and Keller, G. (2004). Development of definitive endoderm from embryonic stem cells in culture. Development 131, 1651–1662.

Li, R., Zhong, C., Yu, Y., Liu, H., Sakurai, M., Yu, L., Min, Z., Shi, L., Wei, Y., Takahashi, Y., et al. (2019). Generation of Blastocyst-like Structures from Mouse Embryonic and Adult Cell Cultures. Cell 179, 687–702 e618.

Martin, B. L. and Kimelman, D. (2008). Regulation of canonical Wnt signaling by Brachyury is essential for posterior mesoderm formation. Dev Cell 15, 121–133.

Moris, N., Anlas, K., van den Brink, S. C., Alemany, A., Schroder, J., Ghimire, S., Balayo, T., van Oudenaarden, A. and Martinez Arias, A. (2020). An in vitro model of early anteroposterior organization during human development. Nature 582, 410–415.

Muzumdar, M. D., Tasic, B., Miyamichi, K., Li, L. and Luo, L. (2007). A global double-fluorescent Cre reporter mouse. Genesis 45, 593–605.

Probst, S., Sagar Tosic, J., Schwan, C., Grun, D. and Arnold, S. J. (2021). Spatiotemporal sequence of mesoderm and endoderm lineage segregation during mouse gastrulation. Development 148.

Rivera-Perez, J. A. and Magnuson, T. (2005). Primitive streak formation in mice is preceded by localized activation of Brachyury and Wnt3. Dev Biol 288, 363–371.

Rossi, G., Broguiere, N., Miyamoto, M., Boni, A., Guiet, R., Girgin, M., Kelly, R. G., Kwon, C. and Lutolf, M. P. (2021). Capturing Cardiogenesis in Gastruloids. Cell Stem Cell 28, 230–240 e236.

Teo, A. K., Arnold, S. J., Trotter, M. W., Brown, S., Ang, L. T., Chng, Z., Robertson, E. J., Dunn, N. R. and Vallier, L. (2011). Pluripotency factors regulate definitive endoderm specification through eomesodermin. Genes Dev 25, 238–250.

Tosic, J., Kim, G. J., Pavlovic, M., Schroder, C. M., Mersiowsky, S. L., Barg, M., Hofherr, A., Probst, S., Kottgen, M., Hein, L., et al. (2019). Eomes and Brachyury control pluripotency exit and germ-layer segregation by changing the chromatin state. Nat Cell Biol 21, 1518–1531.

Turner, D. A., Girgin, M., Alonso-Crisostomo, L., Trivedi, V., Baillie-Johnson, P., Glodowski, C. R., Hayward, P. C., Collignon, J., Gustavsen, C., Serup, P., et al. (2017). Anteroposterior polarity and elongation in the absence of extra-embryonic tissues and of spatially localised signalling in gastruloids: mammalian embryonic organoids. Development 144, 3894–3906.

Turner, D. A., Rue, P., Mackenzie, J. P., Davies, E. and Martinez Arias, A. (2014). Brachyury cooperates with Wnt/beta-catenin signalling to elicit primitive-streak-like behaviour in differentiating mouse embryonic stem cells. BMC Biol 12, 63.

van den Brink, S. C., Alemany, A., van Batenburg, V., Moris, N., Blotenburg, M., Vivie, J., Baillie-Johnson, P., Nichols, J., Sonnen, K. F., Martinez Arias, A., et al. (2020). Single-cell and spatial transcriptomics reveal somitogenesis in gastruloids. Nature 582, 405–409.

van den Brink, S. C., Baillie-Johnson, P., Balayo, T., Hadjantonakis, A. K., Nowotschin, S., Turner, D. A. and Martinez Arias, A. (2014). Symmetry breaking, germ layer specification and axial organisation in aggregates of mouse embryonic stem cells. Development 141, 4231–4242.

van den Brink, S. C. and van Oudenaarden, A. (2021). 3D gastruloids: a novel frontier in stem cell-based in vitro modeling of mammalian gastrulation. Trends Cell Biol 31, 747–759.

Veenvliet, J. V., Bolondi, A., Kretzmer, H., Haut, L., Scholze-Wittler, M., Schifferl, D., Koch, F., Guignard, L., Kumar, A. S., Pustet, M., et al. (2020). Mouse embryonic stem cells self-organize into trunk-like structures with neural tube and somites. Science 370.

Veenvliet, J. V., Lenne, P. F., Turner, D. A., Nachman, I. and Trivedi, V. (2021). Sculpting with stem cells: how models of embryo development take shape. Development 148.

Viotti, M., Nowotschin, S. and Hadjantonakis, A. K. (2014). SOX17 links gut endoderm morphogenesis and germ layer segregation. Nat Cell Biol 16, 1146–1156.

Wilson, V. and Beddington, R. (1997). Expression of T protein in the primitive streak is necessary and sufficient for posterior mesoderm movement and somite differentiation. Dev Biol 192, 45–58.

Wilson, V., Manson, L., Skarnes, W. C. and Beddington, R. S. (1995). The T gene is necessary for normal mesodermal morphogenetic cell movements during gastrulation. Development 121, 877–886.

Wilson, V., Rashbass, P. and Beddington, R. S. (1993). Chimeric analysis of T (Brachyury) gene function. Development 117, 1321–1331.

Wood, H. B. and Episkopou, V. (1999). Comparative expression of the mouse Sox1, Sox2 and Sox3 genes from pre-gastrulation to early somite stages. Mech Dev 86, 197–201.

Wymeersch, F. J., Wilson, V. and Tsakiridis, A. (2021). Understanding axial progenitor biology in vivo and in vitro. Development 148.

Xu, P. F., Borges, R. M., Fillatre, J., de Oliveira-Melo, M., Cheng, T., Thisse, B. and Thisse, C. (2021). Construction of a mammalian embryo model from stem cells organized by a morphogen signalling centre. Nat Commun 12, 3277.

Yamaguchi, T. P., Takada, S., Yoshikawa, Y., Wu, N. and McMahon, A. P. (1999). T (Brachyury) is a direct target of Wnt3a during paraxial mesoderm specification. Genes Dev 13, 3185–3190.

Yu, L., Wei, Y., Duan, J., Schmitz, D. A., Sakurai, M., Wang, L., Wang, K., Zhao, S., Hon, G. C. and Wu, J. (2021). Blastocyst-like structures generated from human pluripotent stem cells. Nature 591, 620–626.

