## Supplementary Figures for "Chimeric 3D-gastruloids – a versatile tool for studies of mammalian peri-gastrulation development"

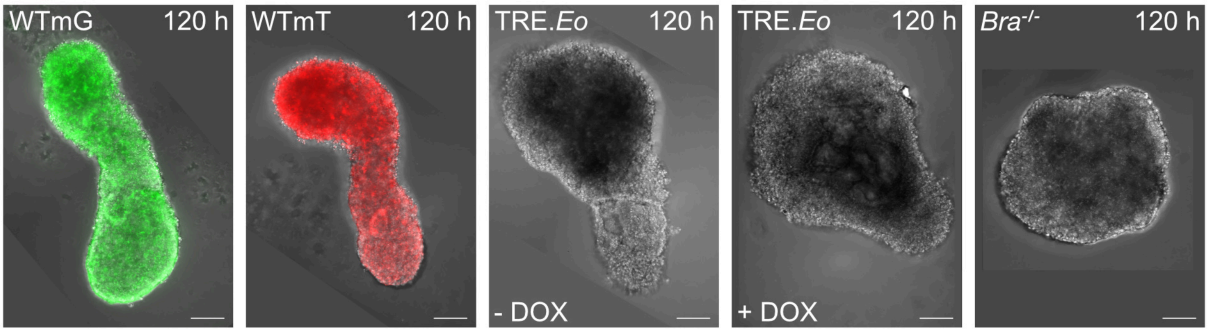


**Supplementary Fig. 1. Baseline analysis of mESC lines used for the generation of chimeric gastruloids.**

Morphological comparison of gastruloids at 120 h generated from cell lines used in this study, WT cells tagged with membraneGFP (WTmG) or membraneTomato (WTmT), WT cells expressing *Eomes* from the dox-inducible promoter (TRE.*Eo*; comparing -DOX vs. +DOX condition), and *Bra*$^{-/-}$ mESCs. Scale bars 100 µm. Experiments were repeated at least twice with at least 48 gastruloids per experiment.

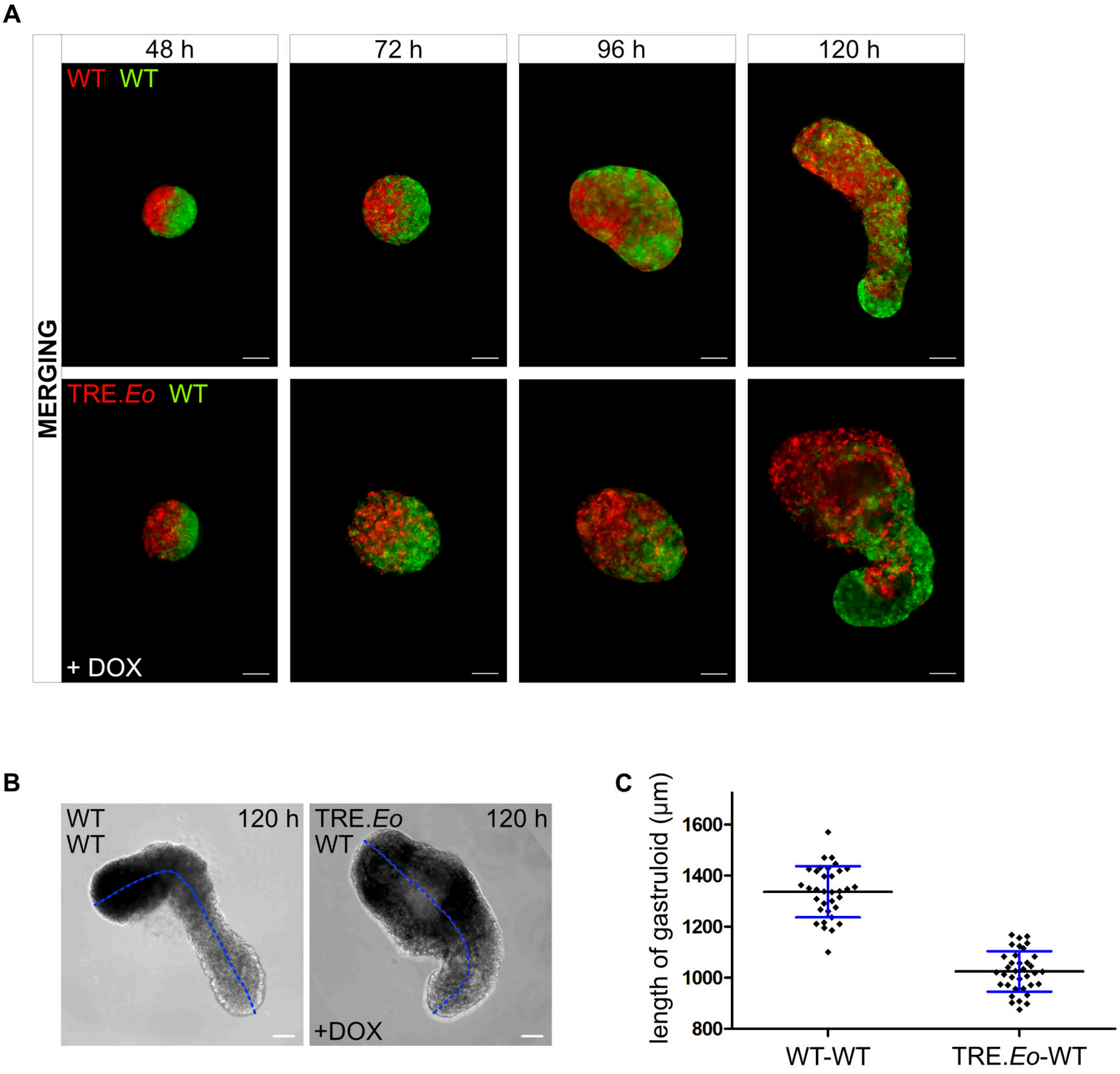


**Supplementary Fig. 2. Induced *Eomes*-expression reduces the axial elongation of merged chimeric gastruloids.**

**(A)** Merged chimeric gastruloids generated by fusing preformed aggregates from WT:WT and TRE.*Eo*:WT cells showing differences in cell distribution following induced *Eomes*-expression (+Dox), analysed at indicated timepoints. Experiments were repeated at least twice with at least 48 gastruloids per experiment.

**(B, C)** Exemplary brightfield images **(B)** and statistical analysis **(C)** of length measurements along the indicated (dashed line) anterior-posterior axis of merged gastruloids for WT-WT (n=34; 1336.5 µm ± 100.3) and TRE.*Eo* -WT cells (n=36; 1024.7 µm ± 79.3) at 120 h, indicating reduced axial elongation of TRE.*Eo*:WT gastruloids. Error bars indicate SD. Scale bars 100 µm.

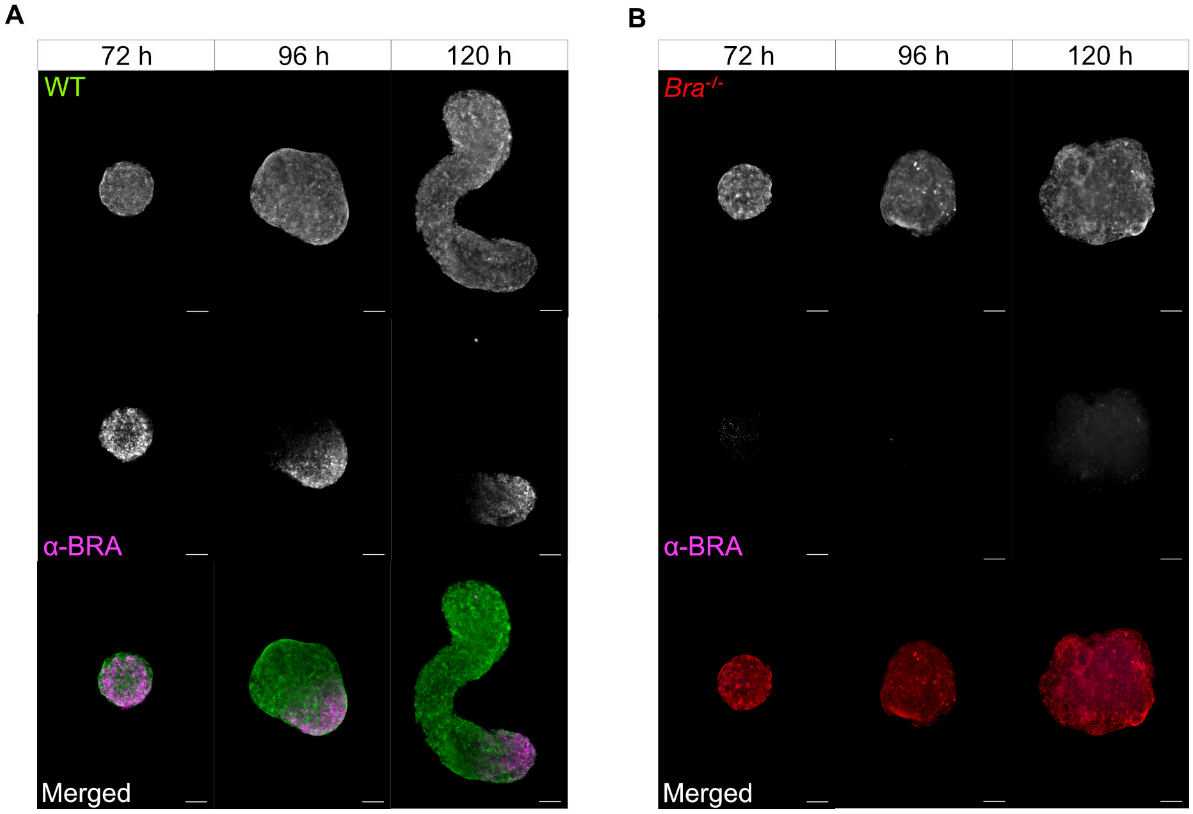


**Supplementary Fig. 3. Baseline analysis of *Bra*$^{-/-}$ mESCs that fail to form elongating gastruloids.**

**(A, B)** Immunofluorescence staining against BRACHYURY (a-BRA) in gastruloids at 72 h, 96 h and 120 h generated from either **(A)** WT, or **(B)** *Bra*$^{-/-}$ mESCs showing the presence of BRACHYURY in the posterior pole of WT, but absence in *Bra*$^{-/-}$ mESC derived gastruloids that fail to elongate. Scale bars 100 µm. Experiment was repeated twice with a minimum of 48 gastruloids per experiment.

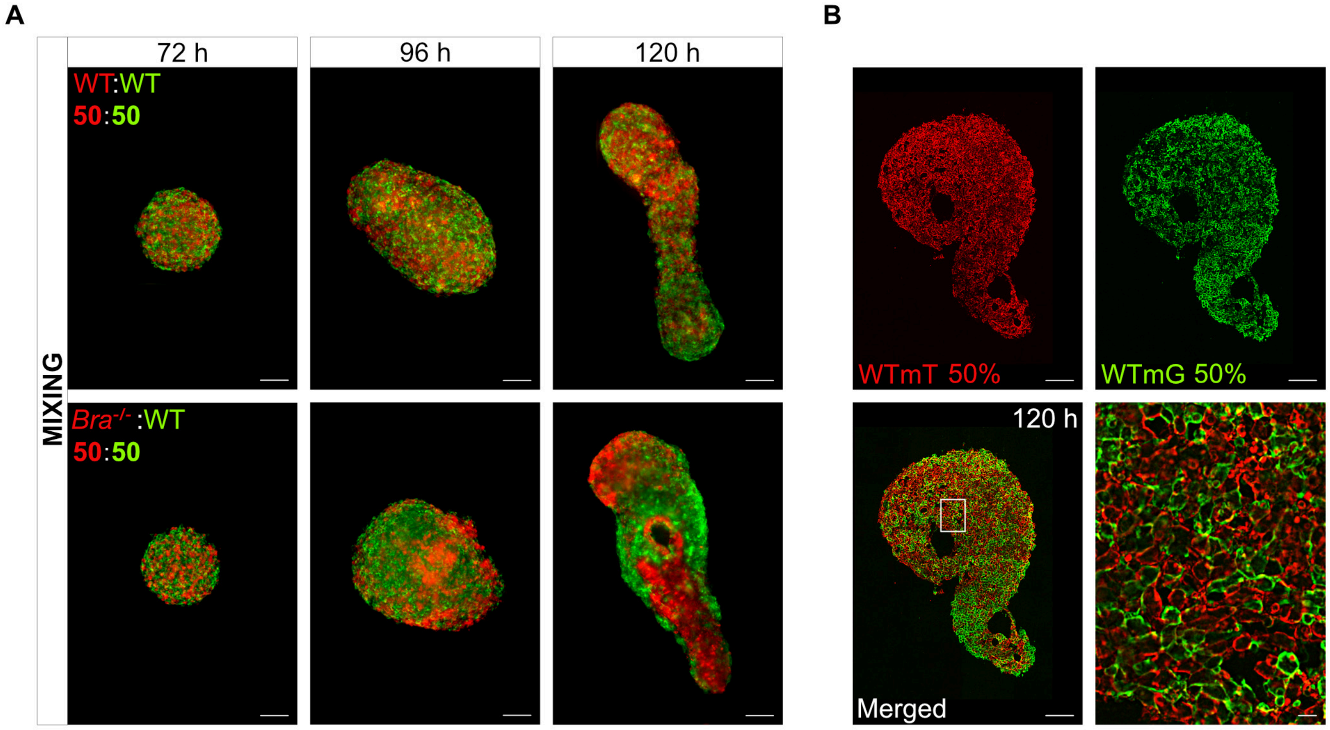


**Supplementary Fig. 4. Comparative analysis of cell sorting behaviour of WT and *Bra$^{-/-}$* mESCs in mixed gastruloids.**

**(A)** Comparative whole mount imaging of cell distribution in 50:50 mixed gastruloids of WT:WT and *Bra$^{-/-}$*:WT cells at 72 h, 96 h, and 120 h, showing differences in cell sorting behaviour between WT and *Bra$^{-/-}$* deficient cells.

**(B)** Histological sections of a gastruloid at 120 h generated by mixing of WTmG and WTmT cells at 50:50 ratio. The high magnification image of the boxed area shows randomly dispersed mT and mG cells without an obvious pattern of cell distribution. Scale bars 100 μm, except 10 μm for the high magnification in (B). Experiments were repeated at least twice with at least 48 gastruloids per experiment.

**Supplementary Movie 1**

**Merging process of preformed ESC aggregates.** A 12 h time-lapse movie showing the merging process of preformed ESC aggregates that adhere quickly and reorganize again into one round aggregate. Scale bar 200 µm.

**Supplementary Movie 2**

**Beating chimeric gastruloid.** A chimeric gastruloid composed of WT and TRE.*Eo* cells at 168 h that displays a beating domain as a result of the induced overexpression of *Eomes*. Scale bar 100 µm.
